# FGMP: assessing fungal genome completeness and gene content

**DOI:** 10.1101/049619

**Authors:** Ousmane H. Cissé, Jason E. Stajich

## Abstract

**Background:** Inexpensive high-throughput DNA sequencing has democratized access to genetic information for most organisms so that research utilizing a genome or transcriptome of an organism is not limited to model systems. However, the quality of the assemblies of sampled genomes can vary greatly which hampers utility for comparisons and meaningful interpretation. The uncertainty of the completeness of a given genome sequence can limit feasibility of asserting patterns of high rates of gene loss reported in many lineages.

**Results:** We propose a computational framework and sequence resource for assessing completeness of fungal genomes called FGMP (Fungal Genome Mapping Project). Our approach is based on evolutionary conserved sets of proteins and DNA elements and is applicable to various types of genomic data. We present a comparison of FGMP and state-of-the-art methods for genome completeness assessment utilizing 246 genome assemblies of fungi. We discuss genome assembly improvements/degradations in 57 cases where assemblies have been updated, as recorded by NCBI assembly archive.

**Conclusion:** FGMP is an accurate tool for quantifying level of completion from fungal genomic data. It is particularly useful for non-model organisms without reference genomes and can be used directly on unassembled reads, which can help reducing genome sequencing costs.

## BACKGROUND

The recent explosion of high-throughput sequencing methods and analytic tools has made sequencing easier and cheaper for nearly all species across the tree of life including uncultivable organisms. However, the quality and completeness of these genomes can vary due to challenges in assembling repeat rich regions and variable or insufficient sequencing coverage (1). Large-scale sequencing projects such as the microbial dark matter project (2), the Human Microbiome Project (3) or the 1000 fungal genomes project (http://1000.fungalgenomes.org) have produced thousands of microbial genome assemblies. The rapid generation and release of draft data is contributing important and useful datasets that are extensively used for studies of pathology, evolution, and discovery of enzymes or pathways. Variable quality and completeness of draft genomes can impact the inferences drawn regarding gene content, transposable element load, and genome size. There is a need to quantify a genome’s completeness to provide context of the quality of information that can be inferred from it. This work is also motivated by observations that lineage specific gene loss is an important driving force in evolution, especially in fungi (4, 5), and the accuracy of conclusions drawn about the patterns of missing genes requires comparisons among similar quality genomes.

Approaches to assess the quality and completeness of a genome have been proposed using nearly 100 different metrics (6). Unfortunately, most of these metrics are generally not applicable to non-model species because they require a substantial amount of additional high-quality data (e.g. fosmids, reference genomes, optical maps) that can be expensive or infeasible to obtain for a large number of samples. Currently, few methods attempt to estimate the amount of missing data in an assembly without prior knowledge. One of the most popular approaches, CEGMA estimates the completeness to the presence of set of 248 single copy gene markers (7, 8). Although CEGMA has been used in numerous studies, a key issue is that makers were selected from only six model eukaryotic species and the ubiquity and detections of these markers may not be consistent as more distant lineages are sampled. CEGMA has been recently discontinued and the authors recommend using alternative tools (http://www.acgt.me/blog/2015/5/18/goodbye-cegma-hello-busco). The concept has been recently revisited and updated with clade-focused sets of protein coding gene markers in BUSCO (9). Another set of 246 single copy fungal gene families has been proposed by FUNYBASE (10). The latter provides a set of conserved fungal genes but the tools are not explicitly developed to assess genome completeness. Furthermore, the FUNYBASE database was generated in 2010 while a broader sampling of diverse fungal genomes is now available (11).

To build a dataset of independent markers to assess completeness, typically, single copy orthologous genes are chosen. Multi-copy gene families are systematically filtered out in these selections, but their utility, as well as that of alternative, non-protein coding gene markers has not been fully explored in assessing genome completeness. Two summary statistics of genome assemblies are frequently used to evaluate quality completeness. The N50 and L50 statistics (12) which describe the level of fragmentation of the assembly are computed based on the lengths of assembly scaffolds or contigs. Both statistics utilize a sorted list of largest to smallest sizes of contigs, where L50 is the length (in bases) of the shortest contig for which 50% of the genome can be contained within contigs of that size or larger, and N50 is the number of contigs that when summed their length is half of the assembly size (13). Note that unfortunately these two concepts are swapped in some tools, where N50 means length and L50 means the count. Still other methods measure the number error per bases or assembly inconsistencies to prediction genome quality (14, 15).

In the present study, we focused on the fungal kingdom. Fungal genome sizes vary from several megabases (Mb) to nearly 1,000 Mb (11). A primary motivation of this work is to provide a realistic estimation of assembly completeness for fungal genomes. The precision dependents on the ability to accurately identify genes, which can appear artifactually fragmented by an incomplete assembly or appear lost due to more rapidly evolving loci in some lineages. The nature, evolutionary trajectory and loss likelihood of genes need to be considered when calculating genome completeness from gene content. We propose a novel set of markers and build a pipeline to assess their presence in genome assemblies called FGMP (Fungal Genome Mapping Project). Our multistep approach extends previous approaches by integrating identifiable fungal protein and highly conserved non-coding regions. The protein markers selected include both single and multi-copy markers and have only a 50% overlap with previously published datasets providing a different dimension of sequence evolution to evaluate the completeness. Highly conserved non-coding regions of fungal genomes are a novel resource we have developed and incorporated into assessment of genome completeness in FGMP. Lastly, we use a multisampling approach coupled to a rarefaction analysis to search for markers in unassembled sequencing reads, which bypass the need for an assembly. Therefore, using FGMP, a researcher can quickly assess the quality of a set of reads in hand before attempting an assembly, which can be computationally expensive. Finally, we described a side-by-side comparison of our tool with state-of arts methods over 246 fungal species with genome assemblies of varying ranges of quality. We captured assembly improvements/degradations in 57 fungal species with more than one released assembly, as recorded in NCBI assembly archive. The modular construction of this work can be a valuable tool for genome completion estimation that can be easily incorporated in more complex pipeline.

## IMPLEMENTATION

A typical run of FGMP consists of three steps. First, a set of raw gene models (proteins) is generated from the queried assembly which are further filtered down to high confidence genes in subsequent steps. Second, the presence of highly conserved non-coding fungal DNA elements (>200 nucleotides) is estimated. Third, the copy number of ubiquitous multi-copy protein families is determined to track possible mis-assemblies or collapsed duplicated regions. The FGMP workflow is diagrammed in Figure 1 and methodology further detailed in the following sections.

**Figure 1.**
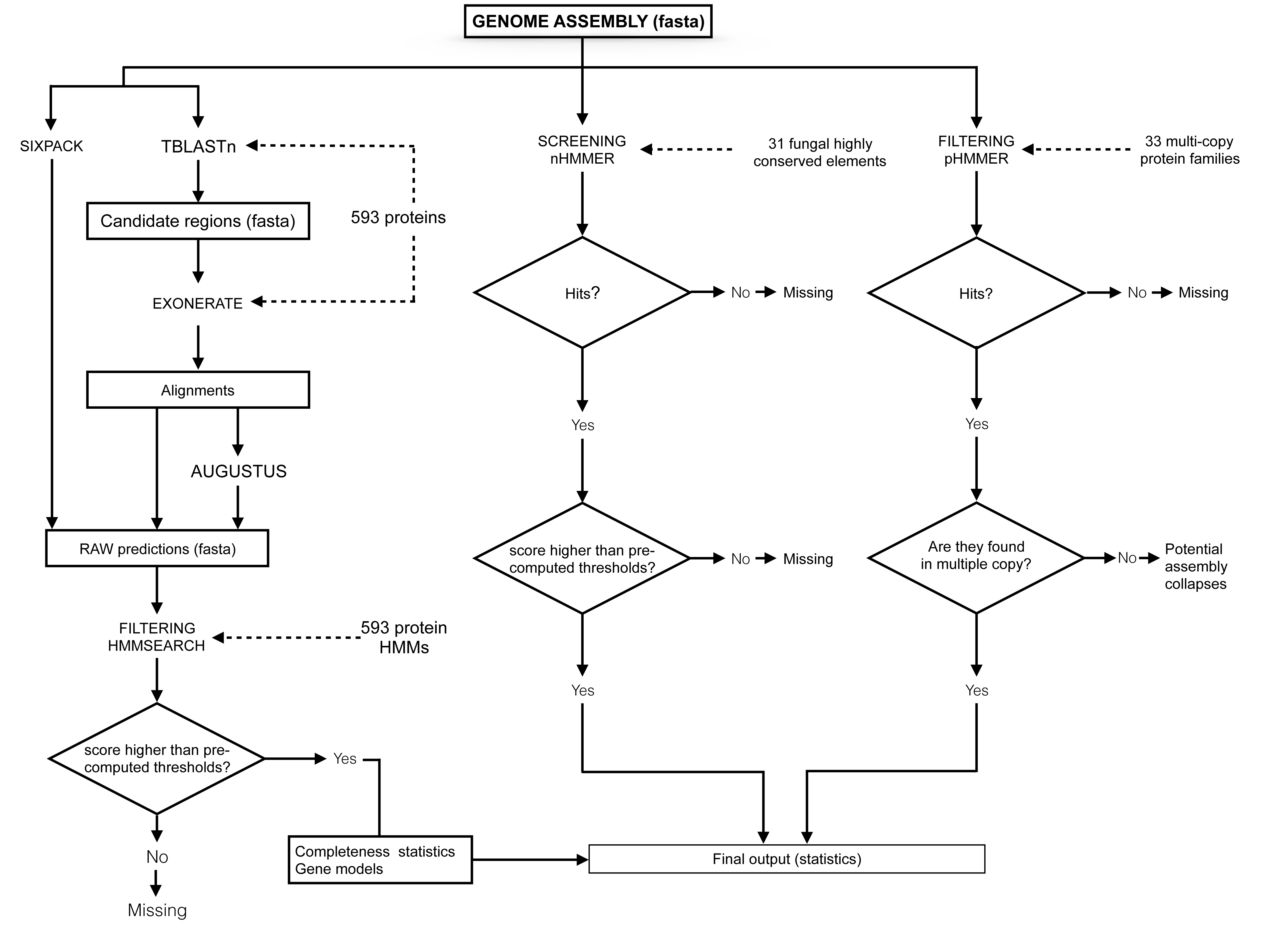
The FGMP workflow. A typical workflow consists of three sequential modules (indicated by the boxes). The first module (FGMP_PROT) automates the use of different programs to evaluate the genome completeness based on pre-defined protein and nucleotide markers. Additional modules evaluate the patterns of conservation fungal multi-copy protein families. FGMP protein and nucleotide datasets are derived from 25 and nine fungal species, respectively (indicated as dotted arrows).

## Reference data preparation

FGMP is primarily designed for assessment of fungal genome quality using defined sets of conserved proteins, noncoding highly conserved DNA elements (HDE) and multi-copy protein families. All the datasets are included in FGMP package and a stable released version is available at DOI: 10.5281/zenodo.1453438. Installation is available via Bioconda package system as “conda install-c bioconda fgmp” (16). Alternatively, a step by step manual installation guide is provided at https://github.com/stajichlab/FGMP.

To generate the protein markers, we analyzed a phylogenomic dataset of 25 fungi with complete genomes covering from all major fungal lineages (Additional file 1). In total, 164,232 proteins were analyzed. Our goal is to capture the fungal protein diversity rather than focusing on universally conserved proteins. We focused on obtaining a set of diverse proteins to be used for initial identification of candidate regions and training sets for gene predictions. Our assumption is that irrespective to the phylogenetic classification of species under analysis, our diversified set of proteins would contain a homolog with sufficient protein similarity to generate a valid gene model. With these concepts in mind, we identified orthologous protein families using OMA (17) followed by inspection of the ortholog clusters using BLAST (18) and full-length pairwise alignments generated using needle from EMBOSS package (19). We extracted 7,773 protein families present in at least four species and use them to construct Hidden Markov Models (HMMs) using HMMER3 (20). In parallel, we selected a single most informative protein in each of these families using M-COFFEE (21), which corresponds to the sequence containing information that is lacking in other sequences of a multiple sequence alignment. We computed the significance scores as follows: each protein of each cluster was compared to its corresponding HMM and the threshold corresponds to 80% of the score of the protein with the lowest score. The use of the full set of 7,773 proteins appeared to be excessively demanding in term of computational resources, even for small sized genome assemblies (e.g. < 8 megabases; data not shown). To reduce the computational burden, we then filtered out potentially paralogous sequences using PHMMER with an E-value of 10^-50^ as cut off (20), and applied the following rules: (i) a marker should be present in at least 99% of the species and (ii) should be unambiguously identifiable based on the alignment score of the protein against the HMM of the family. This filtering reduces our markers dataset to 593 proteins of which 60.3% are from single copy genes contrasting other published strategies, which exclude multi-copy gene families (BUSCO and CEGMA). These 593 representative protein sequences are aligned to the queried genome assembly to identify genomic regions that encode homologous genes using tBLASTn (18). Once candidate regions are narrowed down by these translated alignment searches, fine-grain alignments of the proteins to these homologous regions in the target genome are generated using splice-site aware protein2genome alignment with EXONERATE (22). These alignments-based gene models are used as training sets for AUGUSTUS (23). The predicted proteins including both AUGUSTUS gene models and translated EXONERATE alignment matches are then searched against 593 HMMs to identify the originating genes. FGMP assigns confidence in these predictions based on pre-defined thresholds. We benchmarked FGMP using the full (7,773) and reduced set of proteins (593) on our subsequent analyses and found no significant differences in completeness estimates between the two sets of proteins.

To identify highly conserved non protein-coding fungal DNA elements, we performed pairwise whole genome alignments of nine fungi using LAST (24). The phylogeny of the selected species is presented in the additional file 2. Coding regions were removed from alignments based on NCBI annotations using BEDtools (25). The filtering was carried out using a computational pipeline combining enriched motifs and alignments from MEME (26), BLASTn (18) and EMBOSS ‘needle’ (19). A total of 31 non-coding highly conserved regions in each species were extracted with a requirement that loci be at least 200 nucleotides long with a minimum of 70% global identity. These alignments were converted into HMMs using NHMMER from HMMER version 3.1b2. These genomic segments are specific to fungi because no significant hit was detected to species outside the kingdom using NHMMER with an E-value of 0.1 as cut off against a custom eukaryotic genome database containing *Oomycetes*, *Leishmania* and *Plasmodium* species.

To identify ubiquitous multi-copy protein families within our set of 593 proteins, we surveyed 345 fungal genomes. Thirty-three protein families appear to have more than one copy in all the genomes (HMMSEARCH, E-value < 10^−50^). For each of these 33 proteins, we consider the minimum number of copies. FGMP records the number of copies of these 33 proteins and reports when the copy number is lower than expected.

Lastly, FGMP can estimate the level of completion directly from raw sequences using an iterative reservoir sampling approach. The process starts by splitting the set of reads by chunks of 10^4^ sequences. Then, FGMP randomly selects 1000 chunks using a reservoir sampling approach. This parameter can be modified by the user. Chunks of sequences are iteratively screened for presence of 593 protein makers using BLASTx (24). The number of markers detected is recorded at each iteration. FGMP will stop after 20 successive unsuccessful attempts to detect new markers.

## RESULTS

### Protein markers comparison

To determine if there is overlap among protein markers, we compared FGMP protein markers to proteins used by other tools. Noting that CEGMA has been recently discontinued and FUNYBASE is outdated, this comparison is only for an historical perspective. A total of 7,773 FGMP markers were originally obtained, which was reduced to 593 after the removal of ambiguous markers (see Reference data preparation). We compared the markers selected for FGMP (593 proteins) to those used in CEGMA (248 families, 1,488 proteins), BUSCO fungi (1,438 proteins) and FUNYBASE (246 families, 5,166 proteins). Using reciprocal best BLASTp (E-value < 10^−5^), 49.5% of FGMP protein markers are not found in the other datasets whereas the proportions of unique markers using the same criterion in CEGMA, FUNYBASE and BUSCO are respectively 21.7%, 10.5% and 69.8% (see additional file 3). FGMP proteins tend to be conserved in other eukaryotes but their utility outside the fungal kingdom is not explored in the present study. Transferases and transporters are common (13%). Kinases and helicases are overrepresented in FGMP protein dataset where they represent 10% and 5% of 593 protein markers, respectively as compared to 0.8% and 2% of CEGMA makers; 3.3% and 2% in FUNYBASE markers; 3.3% and 0.7% of BUSCO fungi markers. Kinases and helicases are multi-copy protein families in nearly all fungi and likely this multicopy property is why these genes are not present in other datasets, which actively restrict gene duplicates. Most FGMP kinases have homologs in bacteria and archaea, suggesting that they are ancient. Most of the helicases also have archaeal or bacterial homologs as well and are likely a mix of ancient and derived forms.

### Comparison with related tools

To investigate the ability of different software to detect changes in genome assemblies’ quality, we analyzed the initial and subsequently updated genome assemblies of 45 fungi. As FGMP measures the genome completeness using protein and highly conserved non-coding elements, we refer these modules as FGMP_PROT and FGMP_HCE, respectively. This is because FGMP_PROT is roughly equivalent to BUSCO or CEGMA, but FGMP_HCE is unique to FGMP. FGMP_PROT and FGMP_HCE predict increases in genome completeness in 35% and 31% of the 45 species, respectively whereas BUSCO-fungi predicts increases in 53% and CEGMA in 60% of the species (Figure 2 panels a to d). Overall, the compared methods agree on 16 out of 45 species (Figure 2 panel e). FGMP is the most conservative method at assigning increase in completeness between versions and CEGMA is the most permissive. No statistically significant correlation between FGMP and BUSCO results and various genome statistics was observed (i.e. N50, sequencing coverage, or the sequencing technology used), which is consistent with Assemblathon results (6) showing that completeness metrics are not necessarily correlated among genome assembly statistics. However, CEGMA results appeared to be correlated to the sequencing technology used (Spearman rho = 0.2), that is, assemblies generated exclusively with short reads (e.g. Illumina) tend to have lower rates of increased completeness between versions than those built using long reads (e.g. PacBio). FGMP_PROT and FGMP_HCE results are correlated (Spearman rho = 0.39) but clearly independent, which further highlights the utility of interrogating different genomic regions to assess completeness.

**Figure 2.**
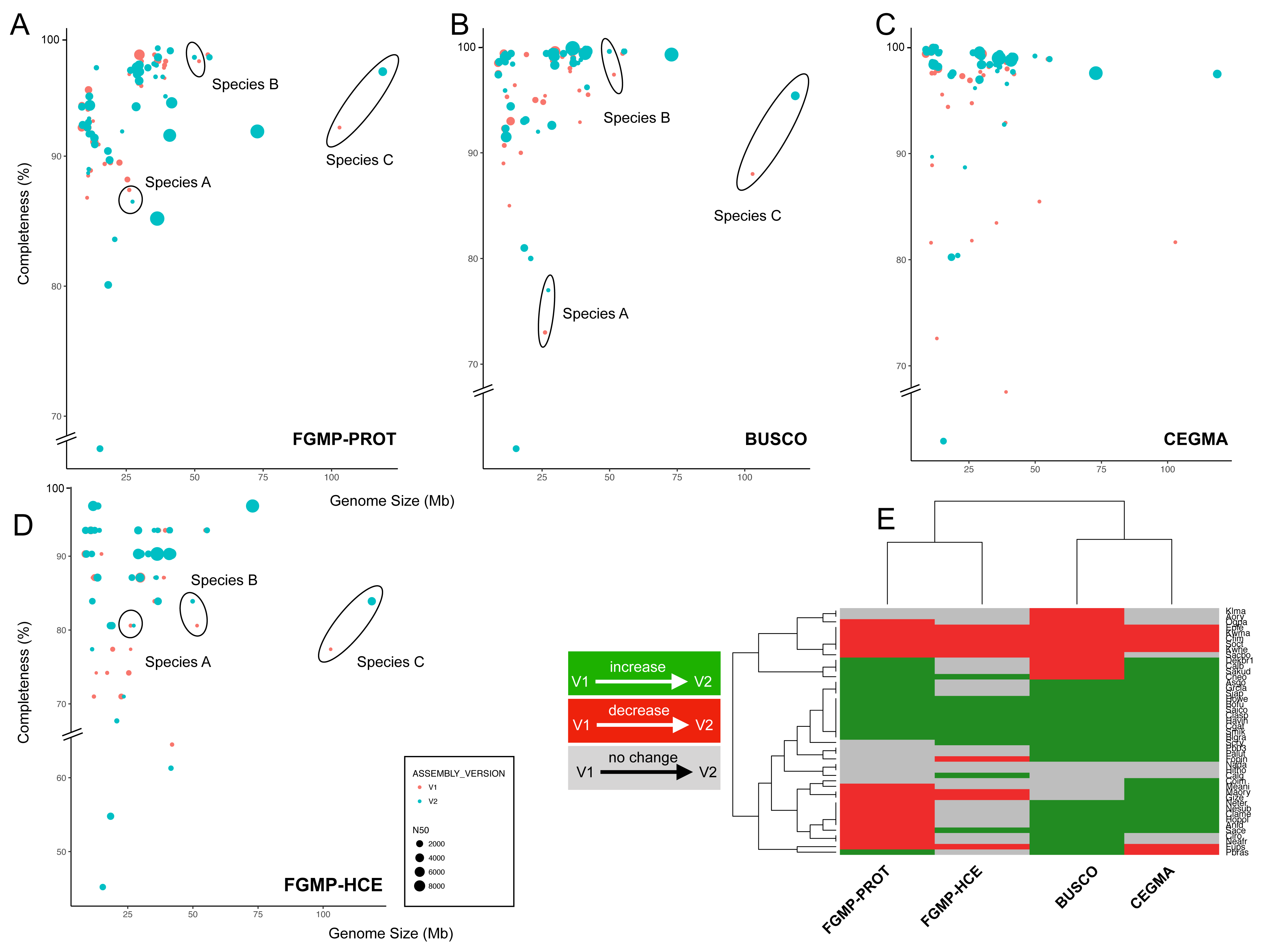
Estimation of genome completeness in fungal genomes. A comparison of genome completeness by multiple software tools using initial and latest genome assembly versions of 45 fungi. Genome completeness expressed as a percentage of expected markers (y-axis) is plotted against assembly size in megabases (x-axis). (A) FGMP completeness estimates based exclusively on protein markers are shown; (B) FGMP estimation based exclusively on highly conserved nucleotide segments; (C,D) BUSCO fungi and CEGMA completeness estimates respectively. In each plot, the dots represent distinct assemblies and color represents their status (red for the initial version and light blue for the latest version), and the diameters are proportional to the N50, a measure of assembly contiguity. Three representative species are highlighted in all panels (ellipses) to show the evolution of genome completeness between assembly versions. (E) Heat map of FGMP, BUSCO and CEGMA completeness estimates.

To further assess the ability of FGMP to detect missing genes and gene loss, we evaluated the impacts of randomly removing ~10% of genomic sequence using 57 fungal genome assemblies (completeness estimates are provided in additional file 4). FGMP_PROT successfully captures the degradations in all assemblies, the average loss rate was estimated at 5% instead of the original 10%, which means the full extent of the simulated loss is not recovered. FGMP_HDE detected the loss of genomic regions in 54 assemblies with an average loss rate of 5.2%. A search with BUSCO (fungi models) captures degradations in 55 assemblies with an average loss rate of 4.3%. The removal of genomic regions prevented CEGMA pipeline from completing in many cases without apparent reasons. Therefore, we have concluded that CEGMA could only be run on the original genome assemblies. Examination of CEGMA completeness estimates found they are not statistically different from BUSCO (fungi models) estimations (Wilcoxon test, *p*- value 0.73), but differ significantly from FGMP results (*p*-value = 0.01). These findings indicate FGMP and BUSCO perform relatively well on genome assemblies of varying degrees of completeness and are similar in estimations detecting degradations in genomes (Figure 3).

**Figure 3.**
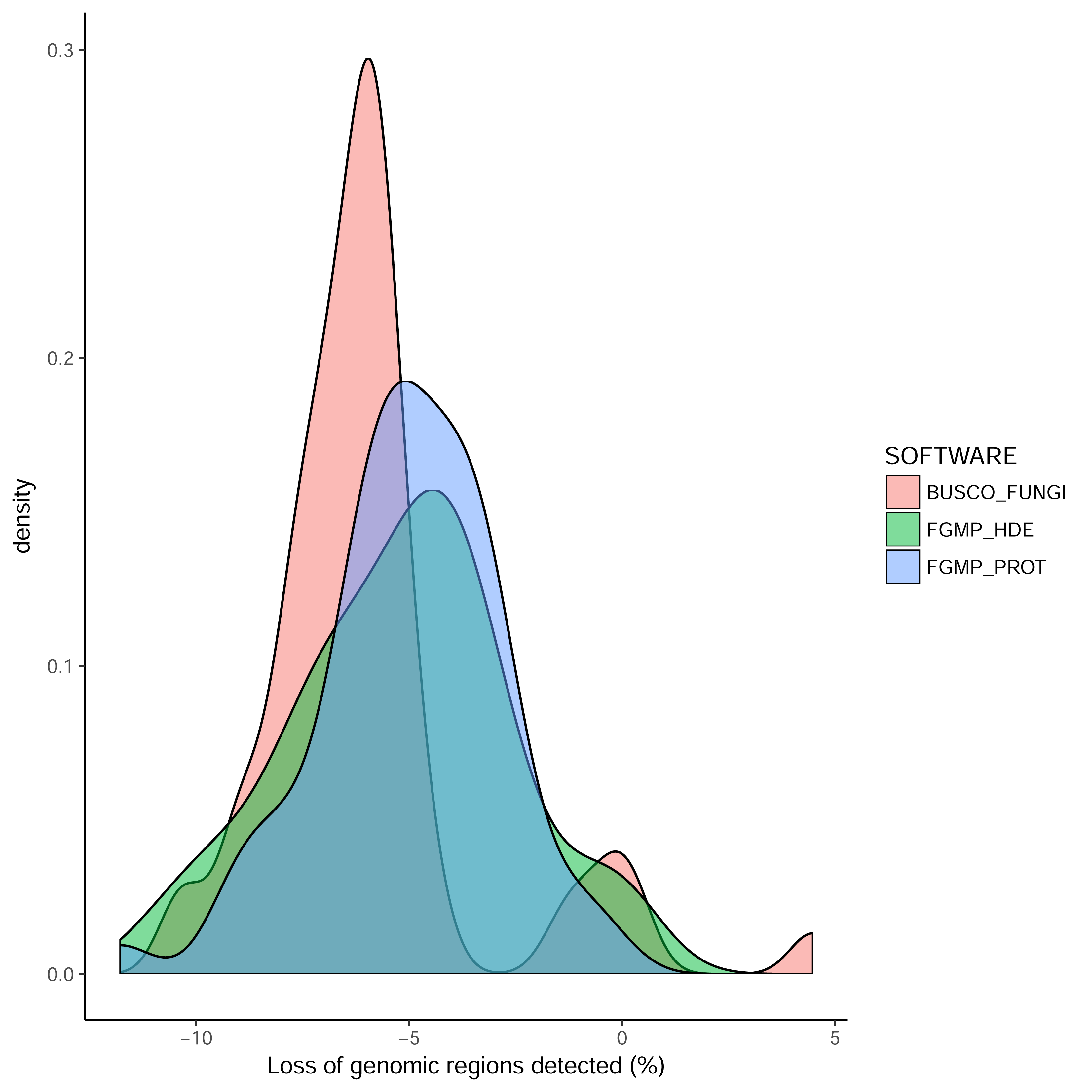
Genomic loss simulations. The genomes of 57 fungal species were randomly truncated to evaluate accuracy of FGMP, BUSCO and CEGMA. The density plots represent the differences (expressed as percentages) between completeness estimates from the full and truncated assemblies.

### Genome completion and ecological or lifestyle traits

We estimated the genome completeness of 166 fungal genomes (species details are described in additional file 7). Only one version of the genome assembly was considered for each species. Each species was classified according to its lifestyle based on published literature (e.g. saprotroph, parasite; references are presented in additional file 7). Parasites are typically characterized by a reduced genome size usually attributed their reliance, either partially or obligately, on nutrients scavenged from hosts. Their genomes are often enriched with transposable and repetitive elements, which in some extreme case composed more than 80% of the genomes (27). Our dataset includes 34 pathogenic species which genome sizes range from 177.6 Mb for the ectomycorrhizal fungus *Cenococcum geophilum* to 2.1 Mb for the microsporidia *Encephalitozoon romaleae*. The remaining 132 fungi were classified as saprotopic and their genome sizes vary from 177 Mb for the ectomycorrhizal fungus *Cenococcum geophilum* to 9.8 Mb for the xerophilic fungus *Wallemia sebi*. Taking the whole set of genomes, the average N50 is 126.7 Mb for an average number of scaffolds per genome of 1,029; an average genome size of 38 Mb and the average fraction of Ns per genome is 3.2%.

Analysis of our set of 166 genome assemblies found that 92% have a CEGMA value > 95% whilst only 58.7% of these assemblies have a 95% completeness with BUSCO fungi, 40% with FGMP_HDE and 54.2% with FGMP_PROT. Genomes labeled as incomplete are typically parasites, which suggest that gene losses from genome streamlining and missing sequence in assemblies might be confounded. Overall completeness predictions correlated with the N50: CEGMA (Spearman rho = 0.35, *P*- value = 1.3 × 10^−8^), BUSCO fungi (R = 0.40; *P* = 2.1 × 10^−11^), FGMP_HDE (R = 0.17; *P* = 0.005) but FGMP_PROT (R = -0.05; *P* = 0.4). FGMP_PROT predictions are not correlated with N50 as this metric incorporates gene fragments, which allow the partial detection of markers even when reliable gene models cannot be built. To avoid overcounting false positives and inflating the estimate, gene fragment sequences are required to score above a predetermined threshold to be accepted as valid hit. However, because short fragments are still required to display a significant similarity versus FGMP protein markers (scores), the likelihood of inflated completeness estimates is expected to be negligible.

### Runtimes

We tracked the running times for 90 fungal genome assemblies (additional file 6). Using six CPU cores each with 8 GB memory (AMD Opteron clock speed 2.1Ggz), FGMP runtimes are proportional to the size and the levels of fragmentation of the genomes under analysis. Runtimes are more influenced by the level of fragmentation of the assembly than its size (additional file 5). For example, FGMP analysis is completed in 39 minutes for the 118 Mb genome size of *B. graminis* (N50 = 2,030. 3 kb) whereas the analysis of the 41 Mb genome of *Magnaporthe oryzae* (N50 = 153 kb) requires three hours (additional file 6). The fastest runtime observed was that of *Cryptococcus gattii* (assembly version 1, size 17.1 Mb, N50 = 44 kb) completed in 22 minutes and the longest with the genome of analysis of the 49.9 Mb genome of *Hortea werneckii* (N50 153 kb) which required four hours.

## DISCUSSION

FGMP is a useful tool for automated assessment of genome assembly completeness of fungal genomes that incorporates measures of gene content covering both protein coding and noncoding regions. The tool combines multilevel analysis by scanning of both coding and non-coding regions of a given genome and provides a detailed reported describing the recovery of multiple types of genomic features in a genome assembly. Compared to existing methods BUSCO and CEGMA, FGMP fills a unique niche by assessing non-coding highly conserved segments and collapsed gene family’s content in addition to measure of protein coding gene conservation. Additionally, FGMP does not rely exclusively on *ab inito* gene predictions with tools like AUGUSTUS which require parameter training. FGMP reports complete, partial and aberrant gene models. FGMP also includes an experimental module, which allow a user to query raw reads using a reservoir sampling approach. This module is currently optimized for low input long reads similar to PacBio or Nanopore sequences. Future versions will include support for estimation from Illumina reads. FGMP has a modular architecture and thus can be easily incorporated into existing genome annotation pipelines.

## CONCLUSION

A realistic estimation of level of genome completeness is a critical metric for accurate comparative genomics studies. This is particularly relevant as the sequencing costs decrease and whole genome assembly is attempted as daily routine for many purposes. BUSCO is currently the only maintained tool for such purpose. FGMP fills a unique niche in the sense that it has modules that assay additional feature types in genomes with no equivalent in existing methods. By applying FGMP to real and simulated datasets, we show that FGMP predictions are reliable and extended that of other software. The tool allows a deeper analysis in the context of evolutionary biology by quickly providing key metrics such the presence of potentially collapsed regions or can be used to screen reads before computationally costly genome assembly is attempted.

## Availability and requirements

Project name: FGMP

Project home page: https://github.com/stajichlab/FGMP

Operating system(s): Linux, Mac OS

Programming language: Perl 5

License: MIT Open Source License

Archived release: DOI: 10.5281/zenodo.1453438

Package system availability: Bioconda

## Availability of data and materials’

<u>Additional file 1:</u> List of fungal species used for phylogenomic analysis.

<u>Additional file 2:</u> Phylogeny of nine fungal species used for the detection of highly conserved nucleotide elements. The divergence times were obtained from http://www.timetree.org(28)

<u>Additional file 3:</u> Comparison of protein markers used for genome completeness estimation.

<u>Additional file 4:</u> Assessment of genome completeness in 57 fungal genome assemblies.

<u>Additional file 5:</u> Scatterplot showing the relationship between FGMP running times and the level of fragmentation for different genome assemblies expressed as N50.

<u>Additional file 6:</u> Genome characteristics, completeness estimates and run times of 90 fungal genomes.

<u>Additional file 7:</u> Lifestyle, genome characteristics and completeness estimates of 166 fungi.

## METHODS

FGMP is written in Perl 5, and is designed for a command line interface. The code is organized into five distinct modules, which are stored in the main library “FGMP.pm”.

1. <u>Identify candidate regions:</u> this module scans the genome assembly using 593 fungal proteins with TBLASTn (18). In parallel, the assembly is translated using SIXPACK (19) and compared to 593 Hidden Markov models of the 593 protein markers using HMMER3 (20). Long FASTA headers are discouraged.
2. <u>Process alignments:</u> aligns 593 protein makers to candidate regions using EXONERATE (22). Alignments are converted in protein sequences and training sets for AUGUSTUS (23).
3. <u>Annotation of candidate regions:</u> uses AUGUSTUS to annotate the candidate regions. The module merges AUGUSTUS predictions with translated proteins from module 2 into a single FASTA file. The module compares raw predictions (proteins or peptides) to 593 HMMs using HMMSEARCH. Lastly, FGMP scans the original assembly for 31 universally conserved fungal elements using NHMMER.
4. <u>Check the status of multi-copy protein families:</u> scans the raw predictions and identify markers that are expected to be in multiple copies. Markers with a lower number of copies than expected are tagged as potentially collapsed regions.
5. <u>Generate final report:</u> gather all raw predictions, filter aberrant predictions (at least twice the average length of the reference makers) and choose the longest gene model for each protein markers.
6. <u>Infer genome completeness from long reads:</u> is triggered when reads are provided. FGMP uses a reservoir sampling approach and BLASTx (18) to search for the 593 proteins markers in the reads.

## Author’s contributions

OHC and JES conceived of the project. OHC wrote the code. OHC and JES unit tested and wrote documentation. OHC generated the data utilized for testing. JES directed the project. OHC and JES wrote the manuscript.

## Acknowledgements

Computations were performed on the University of California-Riverside Institute for Integrative Genome Biology high performance bioinformatics cluster(http://www.bioinformatics.ucr.edu/) supported by NSF MRI DBI 1429826 and NIH S10-OD016290.

